# Engineering synthetic agonists for targeted activation of Notch signaling

**DOI:** 10.1101/2024.08.06.606897

**Authors:** David H. Perez, Daniel Antfolk, Xiomar E. Bustos, Elliot Medina, Shiun Chang, Ahmed A. Ramadan, Paulo C. Rodriguez, David Gonzalez-Perez, Daniel Abate-Daga, Vincent C. Luca

## Abstract

Notch signaling regulates cell fate decisions and has context-dependent tumorigenic or tumor suppressor functions. Although there are several classes of Notch inhibitors, the mechanical force requirement for Notch receptor activation has hindered attempts to generate soluble agonists. To address this problem, we engineered synthetic Notch agonist (SNAG) proteins by tethering affinity-matured Notch ligands to antibodies or cytokines that internalize their targets. This bispecific format enables SNAGs to “pull” on mechanosensitive Notch receptors, triggering their activation in the presence of a desired biomarker. We successfully developed SNAGs targeting six independent surface markers, including the tumor antigens PDL1, CD19, and HER2, and the immunostimulatory receptor CD40. SNAGs targeting CD40 increase expansion of central memory γδ T cells from peripheral blood, highlighting their potential to improve the phenotype and yield of low-abundance T cell subsets. These insights have broad implications for the pharmacological activation of mechanoreceptors and will expand our ability to modulate Notch signaling in biotechnology.

## INTRODUCTION

The Notch pathway is a cell-to-cell communication system that regulates embryonic development, tissue homeostasis, and immune cell differentiation. The signaling of mechanosensitive Notch receptors is tightly regulated^1^ and aberrant Notch activity causes several human diseases^2^. For example, loss-of-function mutations in Notch components are linked to the development of aortic valve disease (Notch1), Alagille syndrome (Notch2, Jagged1), CADASIL (Notch3), spondylocostal dysostosis (DLL3)^3–6^. In cancer, Notch functions as a tumor suppressor or oncogene depending on the cell type, and both loss-of-function and hyperactivating mutations influence tumorigenesis and disease progression^7^. Notch is also pleiotropic with respect to its guidance of cell fate decisions, in that Notch activation stimulates either proliferation or differentiation in different stem cell populations^8^. These diverse functions suggest that Notch agonists and antagonists may each be viable therapeutics in certain biomedical contexts^9^.

The role of Notch in T cell biology has led to the development of several Notch-based strategies for enhancing cancer immunotherapy. Notch signaling is important for several natural stages of T cell maturation^10^, and ex vivo Notch activation is required for the differentiation of T cells from hematopoietic stem cells (HSCs)^11^. This latter function may potentially be used to generate allogeneic T cells for “off-the-shelf” adoptive T cell or chimeric antigen receptor (CAR) T cell therapies. More recently, Notch activation has been shown to enhance the antitumor function of fully mature, activated T cells^12^. Genetic overexpression of an activated form of Notch^13^, as well as culturing T cells in the presence of Notch1-specific antibodies^14^ or ligand-expressing cells^15^, were each associated with improved tumor clearance in various animal models of cancer. Detailed analysis of the T cells used in these studies revealed that these phenotypes were due to the Notch-stimulated induction of exhaustion-resistant or stem-like phenotypes.

At the molecular level, Notch receptors are massive (∼290kD) transmembrane proteins that are activated by a distinctive, mechanical force-driven mechanism^16–18,1^. Notch signaling is initiated when a Delta-like (DLL) or Jagged (JAG) ligand forms a *trans-*interaction with a Notch receptor on the surface of an adjacent cell^1,19–21^. Endocytosis of the ligand then generates a “pulling” force that propagates to the negative regulatory region (NRR) of Notch ^16,22^. This pulling destabilizes the NRR, which exposes internal cleavage sites for processing by the intramembrane proteases ADAM10 (S2 cleavage) and γ-secretase (S3 cleavage)^23,24^. Following these proteolytic events, the Notch intracellular domain (NICD) translocates to the nucleus to function as a transcriptional co-activator^25^.

Although Notch inhibitors are widely available, the requirement for mechanical force in Notch activation has precluded the development of soluble agonists^9,18^. Specifically, these agents are challenging to engineer because they must somehow “pull” on the Notch receptor despite lacking a method of force generation. Several strategies have been developed to activate Notch receptors *in vitro* through mimicry of the physiological activation process. Notch signaling may be induced through co-culture of Notch-expressing cells and ligand-expressing cells, by culturing Notch-expressing cells on plates coated with ligands or antibodies, or by administration of ligand-coated microbeads^14,26,27^. By contrast, only a single antibody targeting Notch3, A13, has been reported to function as a soluble agonist^28^. Binding of this antibody promotes the unfolding of metastable Notch3 NRR domains, which in turn exposes the S2 site for proteolytic cleavage ^29^. Unfortunately, this NRR unfolding approach has been ineffective for receptor subtypes with stable NRRs (e.g., Notch1/2/4), and the lack of soluble agonists remains a significant void in our biochemical toolkit for manipulating the Notch pathway.

In this study, we engineered bispecific proteins that stimulate activation of Notch signaling in desired cellular contexts. We successfully developed SNAGs that enhance the signaling of weakly activating JAG1 ligands, as well as those that selectively activate Notch in the presence of the tumor biomarkers PDL1, CD19, and HER2. Furthermore, we show that SNAGs targeting the immunostimulatory receptor CD40 improve the expansion of γδ T cells from mixed cell populations. The modularity and versatility of this SNAG platform provide a blueprint for the development of a diverse repertoire of Notch-based biologics.

## RESULTS

### Soluble DLL4 ligand multimers do not activate Notch signaling

As an initial attempt to generate Notch agonists, we investigated whether soluble oligomers of an affinity-matured DLL4 ligand (Delta^MAX^) activate Notch signaling^30^. Delta^MAX^ contains ten mutations that increase its affinity for human Notch receptors by 500-to 1000-fold, making it a more potent activator than DLL4 in co-culture and plate-bound formats^30^. We hypothesized that this increased affinity, coupled with receptor crosslinking through multimerization, could introduce tension in the absence of an endocytic pulling force. To test this hypothesis, we incubated Notch1-Gal4 mCitrine reporter cells^31^ with soluble and immobilized Delta^MAX^ multimers (Fig. 1a-c). Delta^MAX^ dimers were generated through the C-terminal addition of a dimeric human IgG1 Fc domain (Fig. 1b), and tetramers were generated by pre-mixing a 4:1 molar ratio of biotinylated Delta^MAX^ with streptavidin (SA, Fig. 1c). We found that neither the monomers nor the multimers induced reporter activity. By contrast, the plated Delta^MAX^ ligands potently stimulated Notch1 activation (Fig. 1a-c). This indicates that the receptor crosslinking by Delta^MAX^-Fc dimers and Delta^MAX^-SA tetramers is insufficient for signaling activation.

**Figure. 1.**
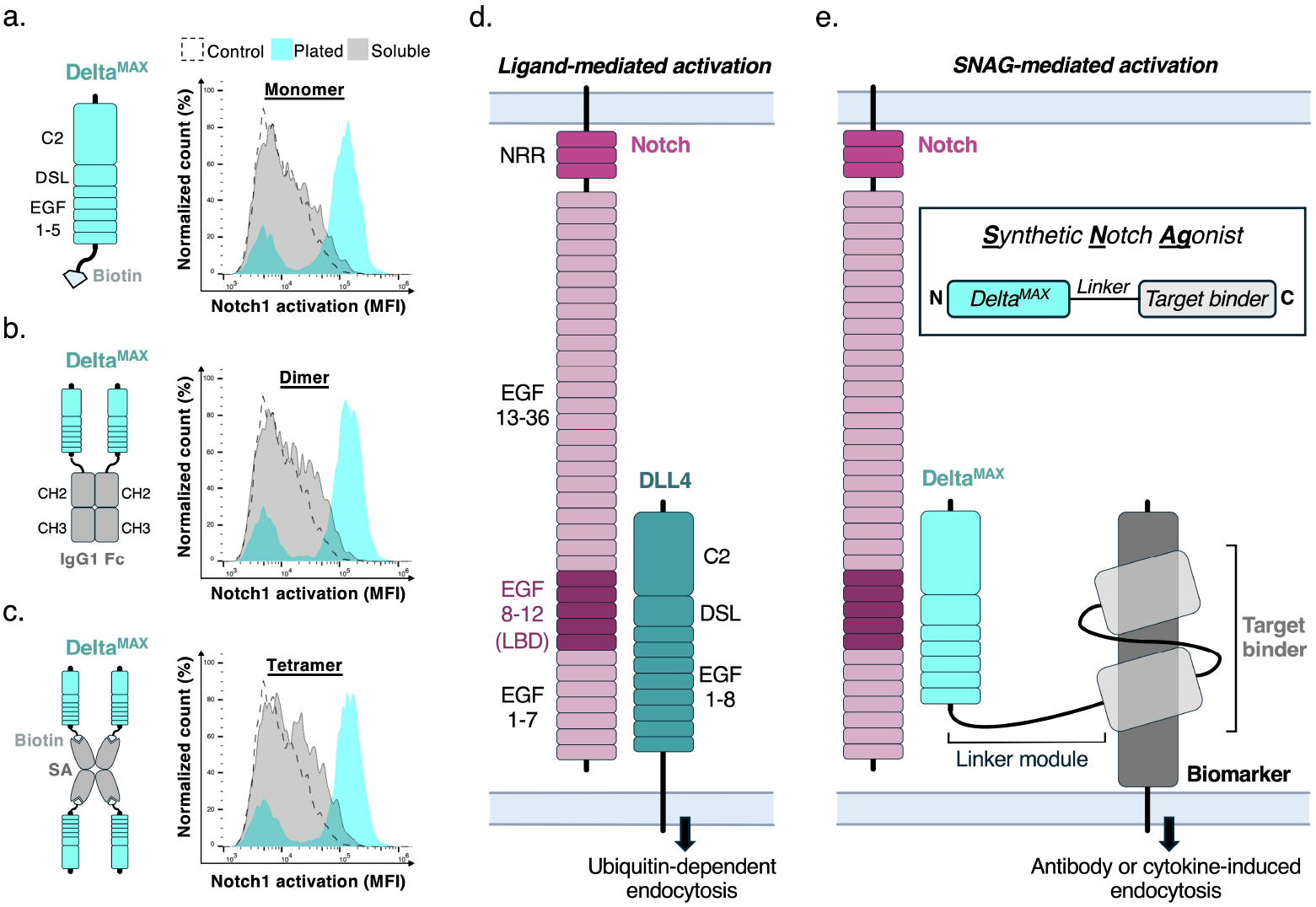
Design concept for synthetic Notch agonists. (a) Flow cytometry histogram overlay of Notch1 reporter cells stimulated by soluble or plated (non-specifically adsorbed) Delta^MAX^. The cartoon depicts the site-specifically biotinylated Delta^MAX^(N-EGF5) construct. (b) Histogram overlay of Notch1 reporter cells stimulated with soluble or plated Delta^MAX^-Fc protein. (c) Histogram overlay of Notch1 reporter cells stimulated with plated or soluble Delta^MAX^-SA tetramers. (d) Cartoon schematic depicting the ECDs of Notch1 and DLL4 interacting during canonical Notch activation. The NRR and ligand-binding domains (LBD) of Notch1 (EGF domains 8-12) are shaded. (e) Schematic of a generalized SNAG construct alongside a cartoon depicting SNAG-mediated Notch activation.

### Design of synthetic Notch agonists

To develop soluble Notch agonists, we engineered bispecific proteins that recapitulate the endocytosis-linked activation mechanism of DLL and JAG ligands (Fig. 1d). SNAGs were created by fusing Delta^MAX^ to the N-terminus of biomarker-targeting antibody fragments via a flexible (GS)5 linker, or by fusing Delta^MAX^ and antibody fragments to the N- and C-termini of a dimeric IgG1 Fc domain (Fig. 1e). These design concepts are intended to form a “molecular bridge” between Notch-expressing cells and cells that express a given surface protein. Conceptually, SNAGs should then activate Notch if the enforced interactions induce endocytic or tensile force capable of unfolding the NRR.

### SNAGs rescue the signaling of a signaling-deficient DLL4 mutant

To demonstrate proof-of-concept, we tested whether SNAGs could rescue the activity of a signaling-deficient DLL4 mutant. Loss-of-function DLL4 cells were generated by expressing a “headless” DLL4 truncation where the Notch-binding C2 and DSL domains^20,32^ were replaced with a BC2 epitope tag (BC2-DLL4^HL^) (Fig. 2a, Fig. S1a)^33^. BC2-SNAGs were then generated by fusing Delta^MAX^ to a BC2-specific nanobody (Figs. 2a-b, Fig. S2a). We found that BC2-DLL4^HL^ cells alone did not activate signaling in a Notch1-Gal4 mCitrine reporter assay, whereas the addition of 1 nM to 100 nM concentrations of SNAGs stimulated a dose-dependent increase in reporter activity (Fig. 2c, Fig. S3a-c). Monomeric BC2-SNAGs containing the (GS)5 linker (BC2-SNAG) stimulated a ∼6-fold increase in Notch1 signaling, whereas dimeric BC2-SNAG Fc fusion proteins (BC2-SNAG^Fc^) were more effective and induced a ∼10-fold increase (Fig. 2c). Importantly, administration of the monomeric or dimeric BC2-SNAGs alone did not substantially increase Notch1 reporter activity, indicating that a mixture of target-expressing and non-expressing cells is required for SNAG-mediated activation (Fig. 2c).

**Figure. 2.**
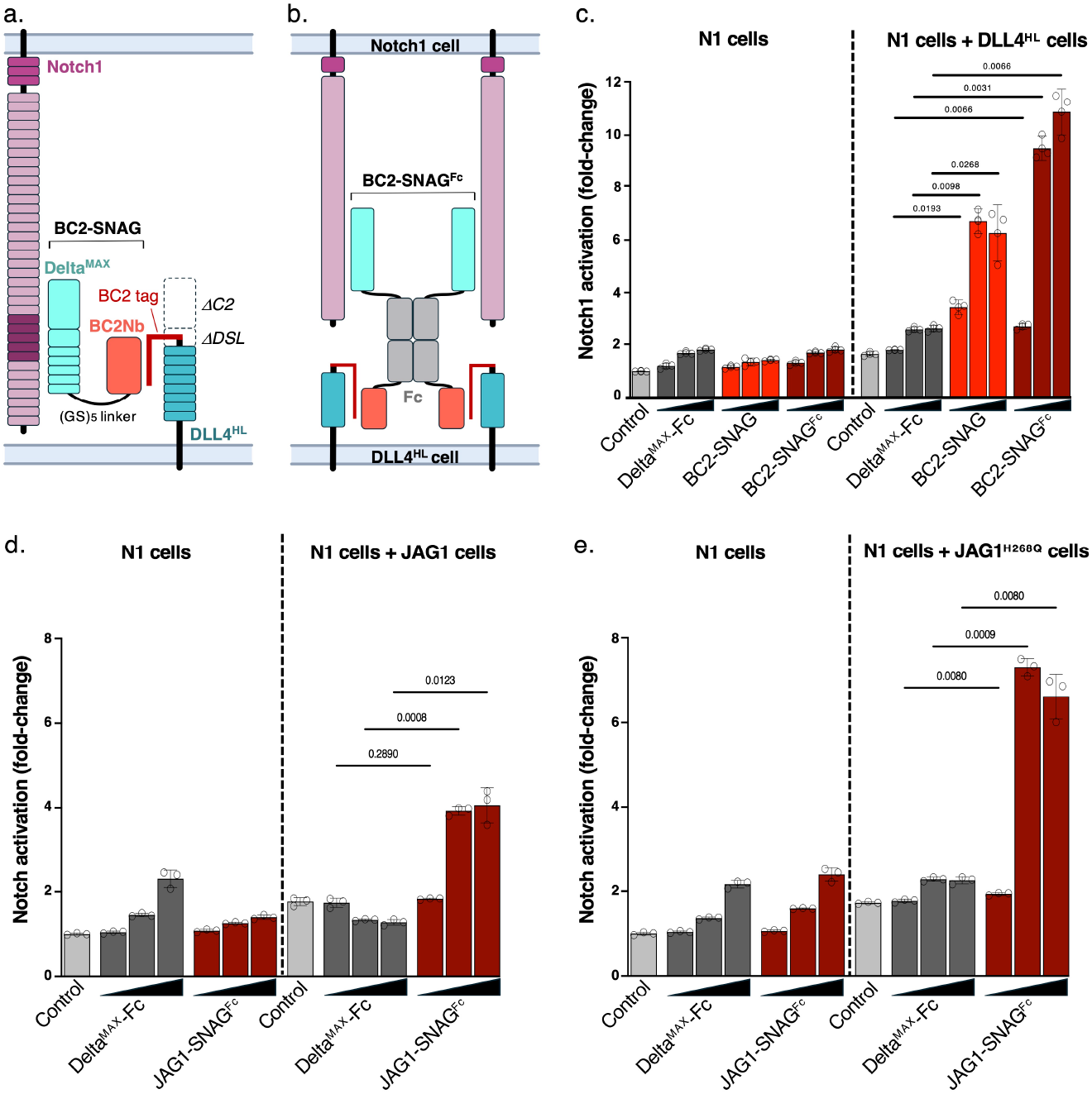
SNAGs rescue the signaling of binding-deficient DLL4 and JAG1 mutants. (a) Cartoon schematic depicting a SNAG binding to Notch1 and a loss-of-function DLL4 mutant. The “headless” loss-of-function DLL4 protein (DLL4^HL^) was generated by replacing the Notch-binding C2-DSL region with a BC2 peptide epitope recognized by the anti-BC2 nanobody. (b) Cartoon schematic depicting the multivalent binding of a dimeric Fc-tagged SNAG (BC2-SNAG^Fc^) to Notch1 and “headless” DLL4. (c-e) Fluorescent reporter assays were used to evaluate SNAG-mediated activation of Notch1 in cocultures with HEK293T cells expressing DLL4^HL^ (c), JAG1 (d), or JAG1^H268Q^ (e). Increasing concentrations (1 nM, 10 nM, or 100 nM) of the indicated SNAGs were added to Notch1-Gal4 Citrine reporter cells alone, or to a 1:1 mixture of Notch1 reporter cells and HEK293 cells expressing DLL4^HL^, and fluorescence was measured by flow cytometry. A representative experiment from three biological replicates is shown. Mean fluorescence intensity (MFI) was normalized to the mean MFI of Notch1 reporter cells alone. Error bars represent the standard deviation of three technical replicates with the *P* value by Student’s *t* test shown above each comparison.

### SNAGs bolster the activity of weakly-signaling JAG1 ligands

DLL or JAG ligands preferentially signal through certain Notch receptor subtypes, and JAG1 is a particularly weak activator of Notch1^34^. Therefore, we tested whether a JAG1-targeting SNAG (JAG1-SNAG^Fc^) could potentiate JAG1-Notch1 signaling. In the JAG1-SNAG^Fc^ construct, Delta^MAX^ and an scFv derived from the JAG1-targeting antibody B70^35^ were fused to the N- and C-termini of an IgG1 Fc domain as described above (Fig. S2b). We then performed a signaling assay to measure the activation of Notch1 reporter cells by JAG1-overexpressing HEK293 cells (JAG1-293 cells) in the presence or absence of the SNAG. We found that addition of the JAG1-SNAG^Fc^ increased Notch1 reporter activity by ∼4-fold compared to JAG1-293 cells alone, and that JAG1-293 cells did not stimulate a significant increase in reporter activity (Fig. 2d, Fig. S1b). We also evaluated the JAG1-SNAG^Fc^ with a JAG1 H268Q “Nodder” mutant that causes Alagille syndrome-like symptoms in mice by decreasing JAG1-Notch1 binding (Fig. S1c). Addition of the JAG1-SNAG^Fc^ to cocultures of Notch1 and JAG1^H268Q^ cells increased JAG1 signaling by up to 7-fold compared to JAG1^H268Q^ cells alone (Fig. 2e). These data indicate that SNAGs can function as “signaling enhancers” by potentiating the activity of endogenous or mutated ligands.

### SNAGs targeting tumor antigens activate Notch in mixed cell populations

We next tested whether SNAGs targeting the tumor antigens PD-L1, CD19, or HER2 (Fig. 3a) can stimulate Notch activation. There is mounting evidence that Notch signaling enhances the function of activated T cells^13–15^, and SNAGs localized to the tumor microenvironment have the potential to stimulate localized activation of tumor-associated lymphocytes. For these SNAGs, the targeting arms were derived from antibody-drug conjugates (ADCs) that were pre-selected for their ability to induce target internalization. We hypothesized that SNAGs incorporating ADC antibodies could thus mimic the physiological endocytosis mechanism of DLL or JAG ligands.

**Figure. 3.**
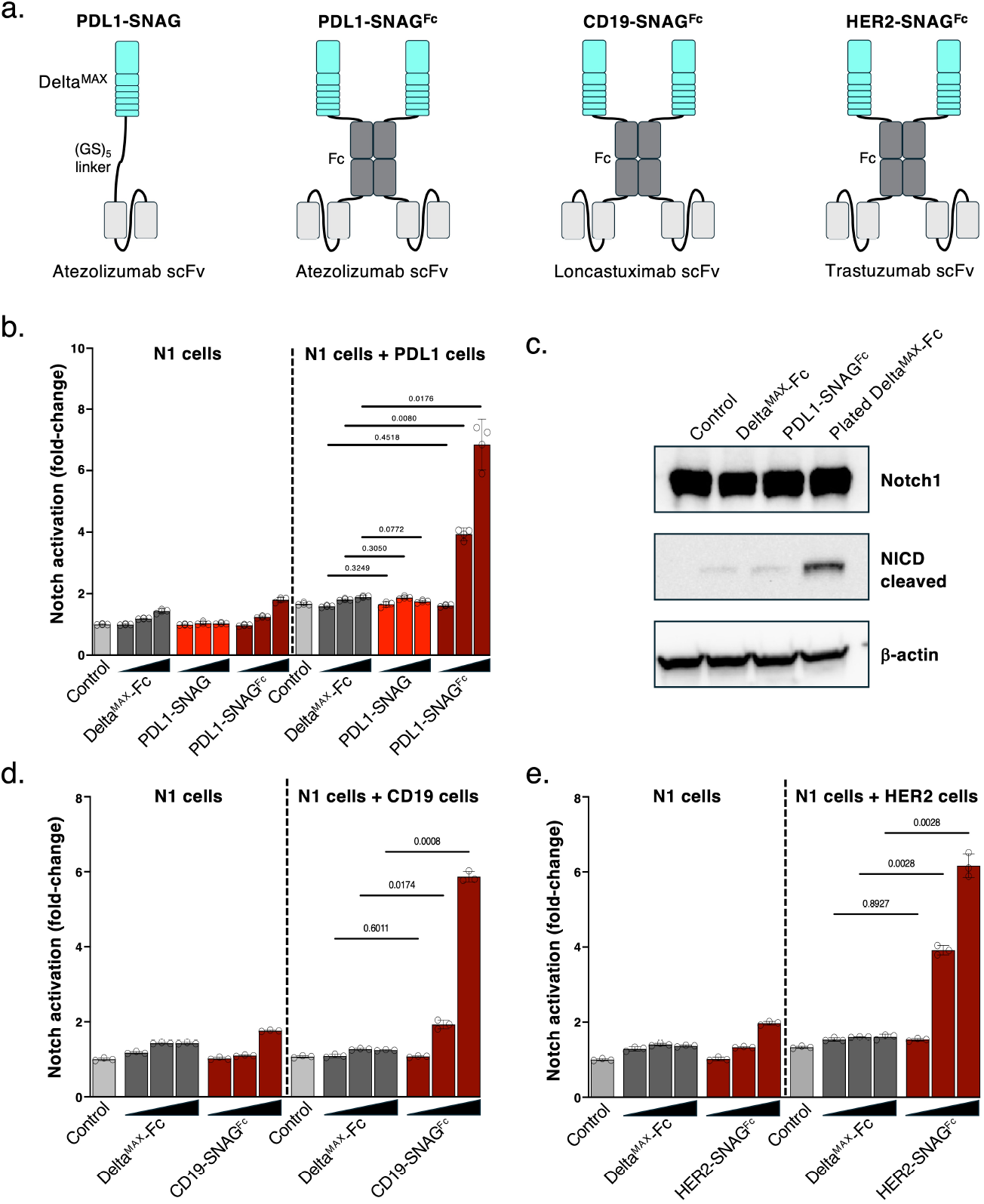
SNAGs targeting tumor antigens activate Notch in mixed cell populations. (a) Cartoon schematics of PDL1, CD19, and HER2 SNAGs. (b) PDL1-SNAG-mediated activation of Notch1 was evaluated in a fluorescent reporter assay. Increasing concentrations (1 nM, 10 nM, or 100 nM) of Delta^MAX^-Fc, PDL1-SNAG, or PDL1-SNAG^Fc^ were added to Notch1-Gal4 mCitrine reporter cells alone, or a 1:1 mixture of Notch1 reporter cells and MDA-MB-231 cells. (c) Activation of Notch1 in MDA-MB-231 cells, which express both PDL1 and Notch1, was assessed by Western Blot using an antibody against the activated NICD. (d, e). Notch1 activation by CD19-SNAGs and HER2-SNAGs, was evaluated using a fluorescent reporter assay. Increasing concentrations (1 nM, 10 nM, or 100 nM) of Delta^MAX^-Fc or each SNAG^Fc^ were added to Notch1-Gal4 mCitrine reporter cells alone, or a 1:1 mixture of Notch1 reporter cells and CD19-overexpressing 3T3 cells (d) or HER2-expressing SK-BR-3 cells (e). For b, d, and e, a representative experiment from three biological replicates is shown. Mean fluorescence intensity (MFI) was normalized to the mean MFI of Notch1 reporter cells alone. Error bars represent the standard deviation of three technical replicates with the *P* value by Student’s *t* test shown above each comparison.

We generated monomeric and dimeric PDL1-SNAGs by fusing Delta^MAX^ to a single-chain variable fragment (scFv) derived from the ADC antibody Atezolizumab^36,37^. In the monomeric PDL1-SNAG, Delta^MAX^ and the scFv were connecting using a (GS)5 linker, and in the dimeric PDL1-SNAG (PDL1-SNAG^Fc^), Delta^MAX^ and the scFv were fused to the N- and C-termini of an IgG1 Fc domain (Fig. 3a, Fig. S2a). Unexpectedly, addition of the monomeric PDL1-SNAG to a 1:1 mixture of Notch1 reporter cells and PDL1-expressing MDA-MB-231 cells did not activate Notch1 (Fig. 3b, Fig. S1d). However, the dimeric PDL1-SNAG^Fc^ protein stimulated a ∼7-fold increase in Notch1 signaling in the coculture, suggesting that multimerization or avidity-enhancement may be required for SNAGs to effectively target biomarkers other than Notch ligands (Fig. 3b). Neither the PDL1-SNAG nor the PDL1-SNAG^Fc^ substantially increased Notch1 reporter activity in the absence of MDA-MB-231 cells. Because of the increased efficacy of the dimeric SNAGs (Fig. 2c, Fig. 3b), we designed subsequent CD19- and HER2-SNAGs using only the Fc-fusion format (Fig. 3a).

### SNAGs do not activate signaling on cells expressing both Notch1 and PD-L1

Given the ubiquitous expression of Notch1 in mammalian cells, it is conceivable that SNAGs could activate signaling when Notch1 and the target protein are both present on the cell surface. To test this possibility, we cultured MDA-MB-231 cells in the presence of soluble Delta^MAX^-Fc, PDL1-SNAG^Fc^, or immobilized Delta^MAX^-Fc and monitored the levels activated Notch1 by Western Blot (Fig. 3c). We found that the plated Delta^MAX^-Fc protein stimulated high levels of Notch1 activation, whereas the PDL1-SNAG^Fc^ did not induce signaling over the background levels observed for soluble Delta^MAX^-Fc alone (Fig. 3c). The inability of SNAGs to activate Notch1 in MDA-MB-231 cells suggests that the present design does not enable sufficient intercellular crosslinking in cultures of cells expressing both Notch1 and the biomarker.

### Development of SNAGs targeting CD19 and HER2

To generate a SNAG targeting the B-lymphocyte antigen CD19, we fused an scFv derived from the CD19-targeting ADC loncastuximab^38^ to the C-terminus of Delta^MAX^-Fc (CD19-SNAG^Fc^, Fig. S2a). The CD19-SNAG was then added to Notch1 reporter cells or to co-cultures of Notch1 reporter cells and CD19-overexpressing 3T3 fibroblast cells (Fig. S1e). We found that the CD19-SNAG^Fc^ protein stimulated up to a 6-fold increase in reporter activity in the co-culture compared to untreated Notch1 cells (Fig. 3d). To generate a SNAG targeting the breast cancer antigen HER2 (HER2-SNAG^Fc^), we replaced the CD19-targeting arm with an scFv derived from the HER2-targeting ADC trastuzumab^39^ (Fig. S2c). Addition of the HER2-SNAG^Fc^ to a mixed culture of Notch1 reporter cells and HER2-expressing SK-BR-3 breast cancer cells induced a 6-fold increase in reporter activity (Fig. 3e, Fig. S1f) at the highest concentration tested (100 nM), which is similar to the level of activation we observed for the PDL1-SNAG^Fc^ and the CD19-SNAG^Fc^ constructs (Fig. 3b, 3d). In the absence of biomarker-expressing cells, neither the CD19-SNAG^Fc^ nor the HER2-SNAG^Fc^ stimulated a significant increase in signaling compared to Delta^MAX^-Fc alone (Fig. 3d-e). Our collective development of PD-L1, CD19, and HER2 SNAGs demonstrates that SNAGs can facilitate signaling by engaging cell surface proteins beyond endogenous ligands.

### Endocytosis is required for SNAG-mediated Notch activation

Ligand endocytosis is important for Notch activation^40^, and this process is regulated by ubiquitination of DLL or JAG ICDs by the E3 ligase Mindbomb1^41–43^. To determine whether endocytosis occurs with a SNAG targeting a surface protein other than a natural Notch ligand, we performed an immunofluorescent endocytosis assay utilizing CD19-SNAG^Fc^ in CD19-expressing cells. CD19-SNAG^Fc^ coupled with a fluorescent secondary antibody (Alexafluor 647-labeled anti-Fc) bound strongly to the surface of the CD19-expressing cells when the mixture was incubated on ice (Fig 4a, Fig. S4a-c), and the contours of the cells were identified by staining for filamentous actin (Fig. S4a). Incubating the cells at 37 °C after attaching CD19-SNAG^Fc^-647 to cells allowed for cellular functions, including endocytosis, to resume. Visualizing the cells after a 15 min incubation at 37 °C showed that the majority of CD19-SNAG^Fc^ is internalized (Fig. 4b, Fig. S4e-d).

**Figure 4.**
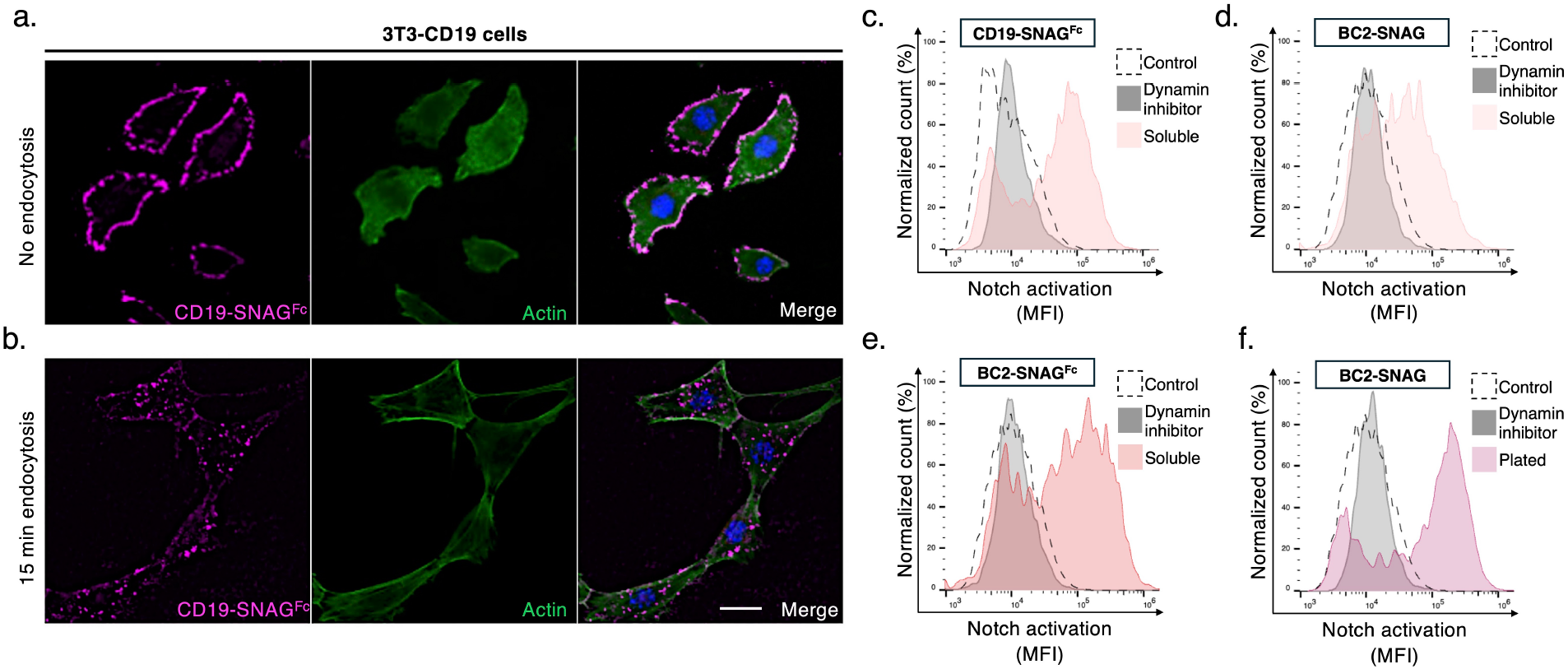
SNAG-mediated Notch activation requires endocytosis. (a) Representative immunofluorescence images of fluorescently labeled CD19-SNAGs (magenta) used to stain the surface of CD19 expressing 3T3 cells that were kept on ice. CD19-SNAGs visualized by Alexa Fluor anti-Fc 647. To visualize the contours of the cells, the actin cytoskeleton was stained using phalloidin-488 (green). Nuclei counterstained by Hoechst 33342 (blue). Scale, 20 mm. (b) Representative immunofluorescence images of fluorescently labeled CD19-SNAGs used to stain the surface of CD19 expressing 3T3 cells, followed by washing away unbound SNAGs and subjecting the cells to a 15 min incubation in a 37 ºC incubator to resume cellular processes including endocytosis. After 15 min the cells were fixed and stained in parallel with the no endocytosis samples. Scale, 20 mm. (d-e) Flow cytometry histogram overlays depicting Notch1 reporter activity induced by soluble CD19-SNAG^Fc^, BC2-SNAG, or BC2-SNAG^Fc^ in the presence or absence of Dynasore. (c) Notch1 reporter cells were co-cultured with CD19-overexpressing 3T3 cells. In (d) and (e), Notch1 reporter cells were co-cultured with HEK293 cells expressing DLL4^HL^. (f) Flow cytometry histogram overlay depicting Notch1 reporter activity induced by immobilized BC2-SNAG in the presence or absence of Dynasore. A representative histogram is shown for each experimental condition from one of three biological replicates.

To test whether endocytosis is necessary for SNAG function, we co-administered SNAGs with the dynamin-dependent endocytosis inhibitor Dynasore. We found that Dynasore completely ablated the activity of CD19-SNAG^Fc^ in co-cultures of Notch1- and CD19-expressing cells, indicating that endocytosis is required for SNAG-mediated activation utilizing CD19 as a biomarker (Fig. 4c). We further found that BC2-SNAGs targeting BC2-DLL4^HL^ were unable to activate Notch1 in co-cultures of Notch1 and BC2-DLL4^HL^ cells in the presence of Dynasore, confirming that endocytosis is also required for SNAG-mediated rescue of DLL4 signaling (Fig 4d-e.) Interestingly, we found that immobilized SNAGs were also unable to activate Notch1 in the presence of Dynasore, suggesting that endocytosis in the Notch-receptor cell is essential for Notch activation by plated ligands (Fig. 4f). These studies demonstrate that Notch activation by plated ligands, SNAGs targeting a DLL4 loss-of-function mutant, and SNAGs targeting tumor antigens each depend on endocytosis. However, it is currently unclear whether endocytosis of the receptor, ligand, or both, is essential for SNAG function.

### SNAGs targeting CD40L increase the expansion of γδ T cells

We next generated a SNAG targeting CD40 (CD40-SNAG), an immunostimulatory receptor that undergoes endocytosis upon binding to CD40 ligand (CD40L)^44^. We hypothesized that a CD40-SNAG could be used to activate Notch in T cells in the presence of CD40^+^ B cells, as these cell types are known to colocalize in germinal centers, peripheral blood, the tumor microenvironment, and other biological contexts^45^. To generate a CD40-SNAG, we fused Delta^MAX^ and the ECD of CD40L to the N- and C-termini of a trimeric leucine zipper^46^ (Fig. 5a, Fig. S2d), respectively. This trimeric scaffold was selected instead of an Fc domain for the CD40-SNAG because CD40L is naturally a homotrimer. We found that addition of the CD40-SNAG robustly activated Notch in mixed cultures of Notch1 reporter cells and CD40-expressing OCI-Ly3 cells (Fig. S1g), but only weakly in Notch1 reporter cells alone (Fig. 5b). Notably, OCI-Ly3 cells used in this assay are non-adherent, indicating that SNAGs may facilitate Notch activation between adherent cells and those grown in suspension.

**Figure 5.**
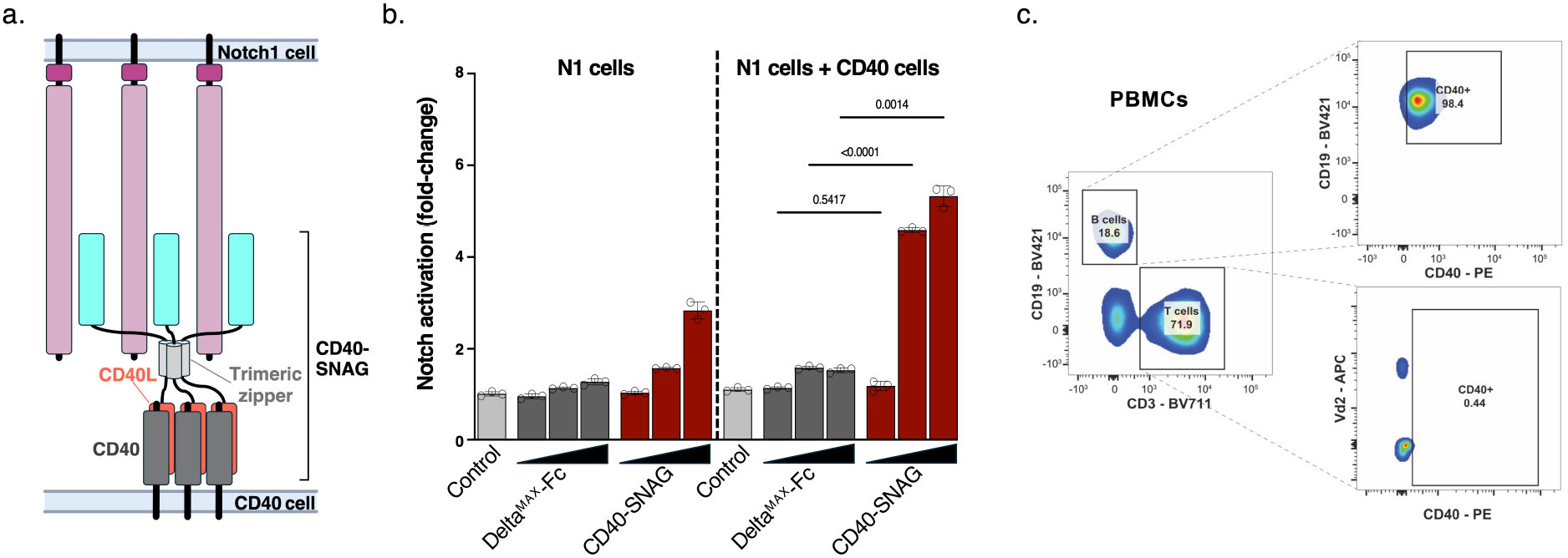
Trimeric CD40-SNAGs activate Notch1 signaling. (a) Cartoon schematic depicting a SNAG binding to Notch1 and CD40 receptor on adjacent cell. The CD40-SNAG was generated by using a GCN4 leucine zipper to trigger dimerization of both the Notch1- and CD40-targeting domains. (b) A fluorescent reporter assay was used to evaluate SNAG-mediated activation of Notch1. Increasing concentrations (1 nM, 10 nM, or 100 nM) of Delta^MAX^-Fc or CD40-SNAG were added to Notch1-Gal4 mCitrine reporter cells alone or a 1:2 mixture of Notch1 reporter cells and OCI-Ly3 cells that have high expression of the CD40 receptor and fluorescence was measured by flow cytometry. A representative experiment from three biological replicates is shown. Mean fluorescence intensity (MFI) was normalized to the mean MFI of Notch1 reporter cells alone. Error bars represent the standard deviation of three technical replicates with the *P* value by Student’s *t* test shown above each comparison. (c) Flow cytometry plots showing the percentage of B cells and T cells on a healthy donor sample, gated on lymphoid, single, live cells. The percentage of CD40+ cells in the B cell subpopulation (CD19+) are shown in the top panel, and the T cell subpopulation (CD3+) on shown in the bottom panel.

There has been a growing interest in γδ T cell-based cancer immunotherapies, including the adoptive transfer of unmodified γδ T cells and the development of γδ T CAR T cells^47–49^. It was previously shown that Notch1 and Notch2 are important for γδ T cell antitumor function^50^, and that Notch stimulation generally induces memory phenotypes in mature and activated T cells^15,14^. Therefore, we sought to determine whether the CD40-SNAG could be used to improve the expansion and phenotype of γδ T cells isolated from peripheral blood mononuclear cells (PBMCs). To determine the ratio of CD40-expressing and non-expressing cells, we analyzed PBMCs by flow cytometry. We found that a representative PBMC culture contained 18.6% CD19+ CD40^+^ B cells and 71.9% CD3^+^ CD40^-^ T cells (Fig. 5c), suggesting that the T cell fraction would be amenable to stimulation with the CD40-SNAG.

To test the effect of the CD40-SNAG on γδ T cells, we administered increasing amounts of CD40-SNAG during an established γδ T cell expansion protocol^51^. Addition of the CD40-SNAG increased both the relative amount and total amount of γδ T cells recovered at all concentrations tested (Fig. 6a-c). We found that the highest CD40-SNAG concentration (500 ng/mL) was associated with >4-fold increase in the total number of γδ T cells compared to untreated PBMCs or those treated with Delta^MAX^-Fc or trimerized CD40L alone. Treatment with 500 ng/mL CD40-SNAG also induced a ∼4-fold increase in the number of central memory cells and a significant reduction in the number of effector memory γδ T cells (Fig. 6d). This bias towards the central memory subset is a desirable outcome given that central memory T cells have increased survival and antitumor function compared with effector memory T cells and effector T cells^52^.

**Figure. 6.**
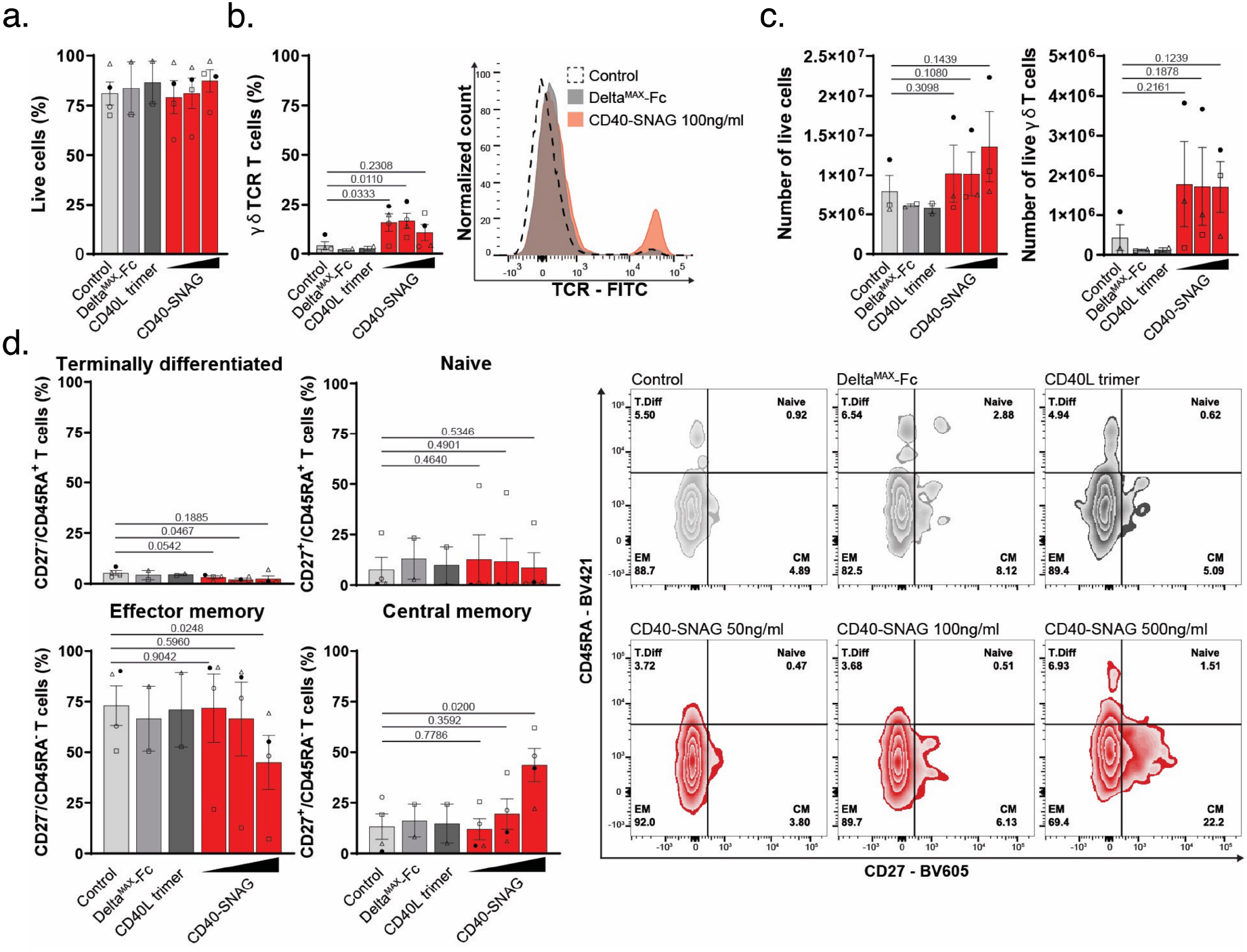
CD40-SNAGs improve the expansion and phenotype of γδ T cells. (a) Bar chart showing the percentage of live cells in culture with different concentrations of CD40-SNAG or controls; gated on lymphoid, single cells. (b) Percentage of γδ TCR expression in T cells cultured with CD40-SNAG vs controls. The histogram to the right shows data from a representative biological replicate; gated on lymphoid, single, viable, CD3+ cells. (c) Bar graph showing the total number of cells in culture by each condition (left) and the relative number of γδ TCR^+^ cells (right). (d) Bar charts representing the distribution of different differentiation phenotypes in γδ TCR^+^ T cells cultured with CD40-SNAG or controls. A representative flow cytometry zebra plot from one healthy donor is shown to the right. Cells were analyzed and counted on day seven, in the bar charts each symbol correspond to an independent donor (biological replicate), p values over the bars were calculated using paired t-tests comparing to the control.

## DISCUSSION

The development of soluble agonists has been an enduring challenge in the Notch field^9,53^. The SNAG platform described here offers a solution to this problem and provides a framework for the development of a diverse array of Notch activating biologics. Such agents have a wide range of potential translational applications, particularly in cancers where Notch functions as a tumor suppressor^7^, T cell manufacturing^11,27^, T cell immunotherapy^13–15^, wound healing^54^, and other areas of regenerative medicine. These first-generation SNAGs were engineered using an Fc-fusion format used in clinically viable protein drugs, which may also help to accelerate *in vivo* translation.

In their present form, SNAGs facilitate potent activation of Notch signaling in mixed populations of cells. However, we anticipate that the design may be tuned to further optimize SNAG function. For example, it may be preferable to engineer SNAGs such that the target binding arm has a higher binding affinity than the Notch-binding arm to improve specificity and tissue distribution. Such strategies have been successfully employed both for bispecific inhibitory antibodies^55^ and T cell engagers^56^. Additionally, higher-order oligomers beyond the dimeric and trimeric SNAG scaffolds tested here may lead to increased signaling potency. Future studies will focus on optimizing affinity and multimerization to maximize signaling while maintaining favorable biochemical properties.

One surprising observation was that PDL1-SNAGs did not activate signaling on cells expressing both PDL1 and Notch1. We speculate that these SNAGs engage the two targets in *cis* on the surface of a single cell, as opposed to bridging PDL1 and Notch1 proteins between cells, and that *cis* interactions do not introduce sufficient tension to unfold the NRR. This may be attributed to the restricted diffusion of SNAGs in the 2-dimensional environment of the membrane, which can promote preferential *cis* interactions by increasing the local concentration. Previous studies have shown that *cis* inhibition of Notch signaling occurs when ligands and receptors are expressed on the same cell^31,57^, and it appears that SNAGs are similarly unable to activate Notch in this context. Regardless, the ability of SNAGs to mediate unidirectional signaling enables highly selective targeting, which could minimize the risks of potential toxicity from global Notch agonism.

There has been a longstanding interest in using Notch stimulation to manufacture “off-the-shelf” CD4^+^ or CD8^+^ T cells for immunotherapy^58^. By contrast, the field of γδ T cell immunotherapy emerged more recently and the role of Notch in their biology is less thoroughly understood^47^. These cells represent only 1-10% of the T cell population in healthy donors, and protocols for the manufacture of pure populations of functional γδ T cells are continuously being optimized^59,60^. We showed that the addition of a CD40-SNAG can greatly increase both the total number of γδ T cells and the fraction of cells with desirable central memory phenotypes. This expansion was not observed upon the addition of trimerized CD40L, suggesting that CD40 signaling alone is not able to recapitulate the effects of the CD40-SNAG. In future studies, it will be interesting to probe whether Notch stimulation in γδ T cells is associated with enhanced antitumor function akin to what is observed for conventional activated T cells^14^, or whether there are synergistic effects between Notch activation in T cells and CD40 activation in B cells.

Although SNAGs are effective in mixed cell populations, the development of “unconditional” agonists that do not rely on a secondary target remains a challenging problem. Thus far, it appears that the metastable NRR of Notch3 is uniquely susceptible to antibody-mediated destabilization^29,61^. The engineering of agonists targeting other Notch receptors with more stable NRRs may require alternative solutions. The successful activation of Notch1 with ligands immobilized on beads^27^ or DNA origami structures^62^ suggests that oligomerization may be an effective strategy, but these methods are not currently viable for in vivo applications. Despite these limitations, the development of SNAGs represents a key first step towards the widespread development of Notch activating molecules for basic and translational research.

## AUTHOR CONTRIBUTIONS

V.C.L. and D.H.P. wrote the manuscript. V.C.L., D.H.P, D.A., and D.A.D. designed the experiments. D.H.P. cloned the SNAG constructs, purified the proteins, and performed the signaling assays. E.M. and D.G.P. generated the Delta^MAX^ constructs. D.A. performed the Notch activation assays in MDA-MB-231 cells and immunofluorescent endocytosis assays. X.E.B. performed T cell expansion assays. P.C.R. and S.C. assisted with immunology experiments. V.C.L. supervised the project and edited the manuscript.

## ACKNOWLEDGEMENTS

This project was supported by NIH R35GM133482 (V.C.L. and D.A.), the Sigrid Juselius Foundation (D.A.) and NIH R35 Diversity Supplement R35GM133482-03S2 (E.M.). V.C.L. is a Rita Allen Scholar. Shared resources were provided by the Moffitt Cancer Center Support Grant NIH P30CA076292.

## MATERIALS AND METHODS

### Protein expression and purification

All SNAG sequences were cloned into a pAcGP67A vector for insect cell production containing an N-terminal gp67 signal peptide and C-terminal 8xHis-tag. Monomeric SNAGs were generated by fusing a truncated version of the Delta^MAX^ protein spanning from the N-terminus to EGF5 (N-EGF5) fused to a biomarker-targeting scFv or nanobody using a flexible (GS)5 linker. Dimeric SNAG^Fc^ constructs were generated by fusing Delta^MAX^ (N-EGF5) and the biomarker targeting module to the N- and C-termini of a human IgG1 Fc domain, respectively. All SNAG^Fc^ constructs contained short GSG-linkers between the Fc sequence and Delta^MAX^ or the targeting module. Published sequences of atezolizumab, trastuzumab, and loncastuximab^63^ were converted into a scFv format prior to being incorporated into SNAGs, and the sequence of the BC2-specific nanobody^33^ was obtained from the Protein Data Bank (PDB ID 5VIN). Each scFv was generated by fusing the C-terminus of the variable heavy (VH) domain to the N-terminus of the variable light (VL) domain with a (GGGGS)3 linker. Biotinylated Delta^MAX^(N-EGF5) protein was generated through enzymatic modification of a C-terminal biotin acceptor peptide (BirA tag) as previously described^30^. The “headless” loss-of-function DLL4^HL^ mutant was generated by replacing the C2 and DSL domains of human DLL4 with the BC2-peptide sequence, which was connected to the N-terminus of EGF1 by a short GSG-linker. The DLL4^HL^ construct was cloned into a pLenti-IRES-Puro vector for mammalian expression.

All SNAG constructs in this study were expressed for by infecting *Trichoplusia ni* insect cell cultures (Expression Systems) at a density of 2 × 10^6^ cells ml^−1^ with recombinant Baculovirus. Culture supernatants were harvested after 48h, and proteins were purified by nickel and size-exclusion chromatography. Biotinylated proteins were site-specifically modified using BirA ligase and excess biotin was removed by purifying the proteins on a size-exclusion column. Protein purity was assessed by SDS–PAGE using TGX 12% Precast gels (Bio-Rad). All proteins were flash-frozen in liquid nitrogen and stored at −80 °C following purification.

### Cell culture and generation of cell lines

Mammalian cells were cultured at 37 °C, with a humidified atmosphere of 5% CO2, washed with Dulbecco’s PBS (DPBS, Corning), and detached with trypsin–EDTA 0.25% (Gibco) for subculturing or cell-based assays. Notch reporter cell lines CHO-K1 N1-Gal4 were a gift from Dr. Michael Elowitz (California Institute of Technology)^31^. Briefly, transfections of HEK293T cells were carried out with packaging vectors VSV-G and d8.9 in the presence of polyethyleneimine at a ratio of 4:1 (DNA:polyethyleneimine). HER2^+^ SK-BR-3 cells, human CD19-overexpressing 3T3 cells, PD-L1^+^ MDA-MB-231, and CD40^+^ OCI-Ly3 cells were gifts from Drs. Brian Czerniecki, Fred Locke, Eric Lau, and John Cleveland, respectively (Moffit Cancer Center). HEK293T, SK-BR-3, 3T3 mouse fibroblast, and MDA-MB-231 cells were cultured in high-glucose DMEM (Cytiva) supplemented with 10% FBS (peak serum) and 2% penicillin/streptomycin (Gibco). Puromycin 5 μg ml^−1^ was added to HEK293T cell cultures to maintain homogeneous populations of receptor-expressing cells. CHO-K1 N1-Gal4 cells were cultured in minimum essential medium Eagle-alpha modification (α-MEM, Cytiva) supplemented with 10% FBS (peak serum), 2% penicillin/streptomycin (Gibco), 400 μg ml−1 of zeocin (Alfa aesar) and 600 μg ml−1 of geneticin (Gibco). Expression of receptors on the cell surface was confirmed by flow cytometry (BD Accuri C6 plus) staining the cell lines with anti-hDLL4 PE, anti-JAG1 APC, anti-hPDL1 FITC, anti-hHER2 (anti-IgG FITC), anti-hCD19 FITC, and anti-CD40 PE in DMEM supplemented with 10% FBS for 1 h at 4 °C.

### Notch activation with Delta^MAX^ multimers

On day one, biotinylated Delta^MAX^, Delta^MAX^ tetramers formed with streptavidin, or Delta^MAX^-Fc were reconstituted in DPBS and adsorbed to tissue culture 96-well plates (Costar) for 1 h at 37 °C. The wells were then washed three times with 200 μl of DPBS to remove unbound proteins. Next, CHO-K1 N1-Gal4 cells were detached with trypsin–EDTA 0.25% (Gibco) and manually counted. Appropriate dilutions were prepared in α-MEM media to ensure 30,000 CHO-K1 N1-Gal4 cells per well in a volume of 50 mL. Cells were transferred to the ligand-coated plates and cultured for 24 h at 37 °C in 5% CO2. On day two, CHO-K1 N1-Gal4 cells were washed with 200 μl DPBS, detached with 30 μL of trypsin–EDTA 0.25%, and quenched with 170 mL of α-MEM media. Finally, cells were resuspended, and the H2B-mCitrine signal was measured by flow cytometry (BD Accuri C6 plus). CHO-K1 N1-Gal4 cells alone were used as the control. The measurements represent the mean fluorescent intensity as fold-change of Notch activation ± s.d. of three technical replicates. Notch activation was normalized to wells containing CHO-K1 N1-Gal4 cells alone.

### Notch activation with SNAGs in coculture of cells expressing the target tumor biomarker

On day one, cells expressing the target receptor of the SNAG (signal-sending cells) were detached with trypsin–EDTA, counted manually, and dilutions prepared such that 50 mL of DMEM containing 15,000 signal-sender cells were added to wells of a tissue culture 96-well plate. The next day, CHO-K1 N1-Gal4 reporter cells (signal-receiver cells) were detached with trypsin– EDTA, and 50 μl of α-MEM media containing 30,000 cells were added to the tissue culture 96-well plate containing the signal-sending cells after combining with the indicated Delta^MAX^ or SNAG protein. For the CD40-SNAG experiments, the difference was that an equal amount of signal-sending and signal-receiving cells were added same day. Wells without signal-sending cells were used to determine background activation of Notch by Delta^MAX^ and SNAGs. When testing inhibition of endocytosis, 80 μM of the Dynamin inhibitor I (Dynasore, Sigma) was added to the mixture of Notch reporter cells with protein and added to the tissue culture 96-well plate containing the signal-sending cells. Notch activation was measured as previously described.

### Western blot detection of Notch1 activation by the PDL1-SNAG^Fc^ in MDA-MB-231 cells

Delta^MAX^ (100 nM protein in 600 mL of DPBS) was non-specifically adsorbed to a single well of a 12-well plate for 1 hour at 37 ºC as a positive control for Notch1 activation. The positive control well and three additional wells were then seeded with 200 × 10^3^ cells with MDA-MB-231 cells. The plate was centrifuged at 400 x *g* for 4 min to ensure cells were retained at the bottom of each well, and then the media of all wells was discarded. In the first uncoated well, 600 mL of DMEM was added as a negative control. The second well was filled with 600 mL of media containing 100 nM of Delta^MAX^-Fc to monitor Notch1 activation by soluble ligand. The third was filled with 600 mL of media containing 100 nM PDL1-SNAG^Fc^. The following day, the media was aspirated from all four wells, and the samples were resuspended in 60 mL of Laemli sample buffer with 5% b-mercaptoethanol to lyse cells, followed by boiling at 100 °C for 4 min. Lastly, the samples were analyzed by western blotting using equal protein amounts of cell lysates separated by SDS–PAGE (12% Mini-PROTEAN TGX Precast Protein Gels, Bio-Rad) and transferred to PVDF membranes using an iBlot2 Gel Transfer Device (Thermo Fisher Scientific). The membranes were blocked in 3% BSA + 0.1% TBS-Tween. Primary antibodies were anti-Notch1 (D1E11 rabbit mAb, Cell Signaling Technology, 1:1,000), anti-cleaved Notch1 (Val1744 rabbit mAb, Cell Signaling Technology, 1:1,000), and b-actin (rabbit polyclonal Ab, Cell Signaling Technology, 1:1,000). Secondary antibody anti-Rabbit IgG conjugated to HRP (Goat polyclonal Ab, Vector Laboratories, 1:8,000) was used for detection of proteins using SuperSignal West Pico PLUS Chemiluminescent Substrate (Thermo Fisher Scientific). Images were acquired using a Chemidoc Imaging System and analyzed with Image-Lab v.6 software (Bio-Rad).

### Immunofluorescent cell staining

For endocytosis assays, cells were grown on glass-like polymer bottoms in 24 well black frame plates (Cellvis). For visualization of CD19-SNAG^Fc^ protein binding, 500 nM protein was preincubated with anti-Fc 647 (Alexa Fluor) at 1:200 dilution for 1h on rotation in +4°C. The CD19-SNAG^Fc^-647 solution was added to cells on ice that were further kept in +4°C for 1 h. For endocytosis, the incubation was followed by washing away non-bound CD19-SNAG^Fc^-647 with PBS, and 37°C DMEM added to the cells followed by a 15 min incubation in a 37°C incubator. After incubation of CD19-SNAG^Fc^-647 with or without endocytosis, the cells were fixed in 3% paraformaldehyde and permeabilized with 0.15% Triton X-100 in PBS for 10 min at RT. Nonspecific binding was blocked by incubation in 3% BSA in PBS with 0.05% Triton X-100 and 0.1M glycine for 60 min at RT. Cells were further stained for filamentous actin with Alexa 488 conjugated to phalloidin (Invitrogen) for 45 min to visualize contours of the individual cells. Hoechst 33342 (Invitrogen) was used to counterstain nuclei. Images were acquired using a Keyence BZ-X710 microscope using a Nikon Plan Apo 20x objective. The far-red channel (magenta) was processed with the de-haze function in the BZ-X710LE analyzer software. A minimum of 100 cells were imaged for each condition.

### γδ expansion from PBMCs with CD40-SNAG

Peripheral blood mononuclear cells (PBMCs) were isolated from healthy-donor buffy coats using an established density gradient protocol^51^. γδ expansion was performed using zoledronic acid in addition or not of CD40-SNAG. Briefly, ten million cells per condition were resuspended in RPMI [supplemented with 5%FBS, antibiotic, IL-2 (100 IU/ml) and zoledronic acid (4μM)] at 1×106 cells/ml. The cells were spiked with CD40-SNAG at three different concentrations (50, 100 or 500ng/ml), Delta^MAX^-Fc (100ng/ml), or CD40L trimer (100ng/ml), and cultured in 24-well plates (2ml/well) for three days at 37C (5% CO2). On day three, the full media was replaced by new media with zoledronic acid and CD40-SNAG, Delta^MAX^-Fc, or CD40L trimer as on day one. Two days later, cells were lifted from the plates, centrifuged, and resuspended in new RMPI media [5%FBS, antibiotic, IL-2 (100 IU/ml)] without additional molecules. At day seven, T cells were counted, and their phenotype was assessed by surface staining and flow cytometry (using markers for human anti-CD3 – BV711, anti-Vd2 TCR – FITC, anti-CD45RA – BV421, anti-CD27 – BV605, and aqua live/dead fluorescent dye).

