## Supplemental Figures & Legends for "Engineering synthetic agonists for targeted activation of Notch signaling"

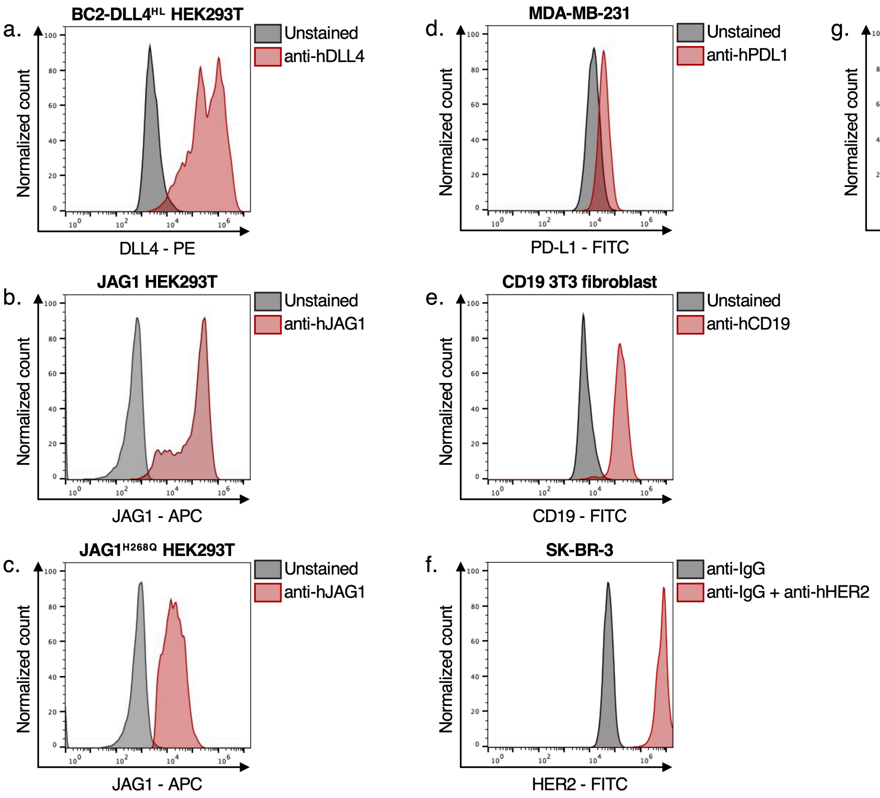


**Figure S1. Expression of SNAG target biomarkers on various cell lines.** Cell lines used in the study were analyzed by flow cytometry to measure surface expression of the indicated biomarker. In each panel (a-f), flow cytometry histograms show the staining of each cell line with the indicated antibody. (a) BC2-DLL4$^{HL}$ HEK293T cells were stained with a fluorescently labeled anti-DLL4 antibody, (b-c) JAG1 HEK293T cells or JAG1$^{H268Q}$ HEK293T cells were stained with an anti-JAG1 antibody, (d) MDA-MB-231 cells were stained with an anti-PDL1 antibody, (e) CD19-expressing 3T3 cells were stained with an anti-CD19 antibody, (f) SK-BR-3 cells were stained with an anti-HER2 antibody, and (g) OCI-Ly3 cells were stained with an anti-CD40 antibody.

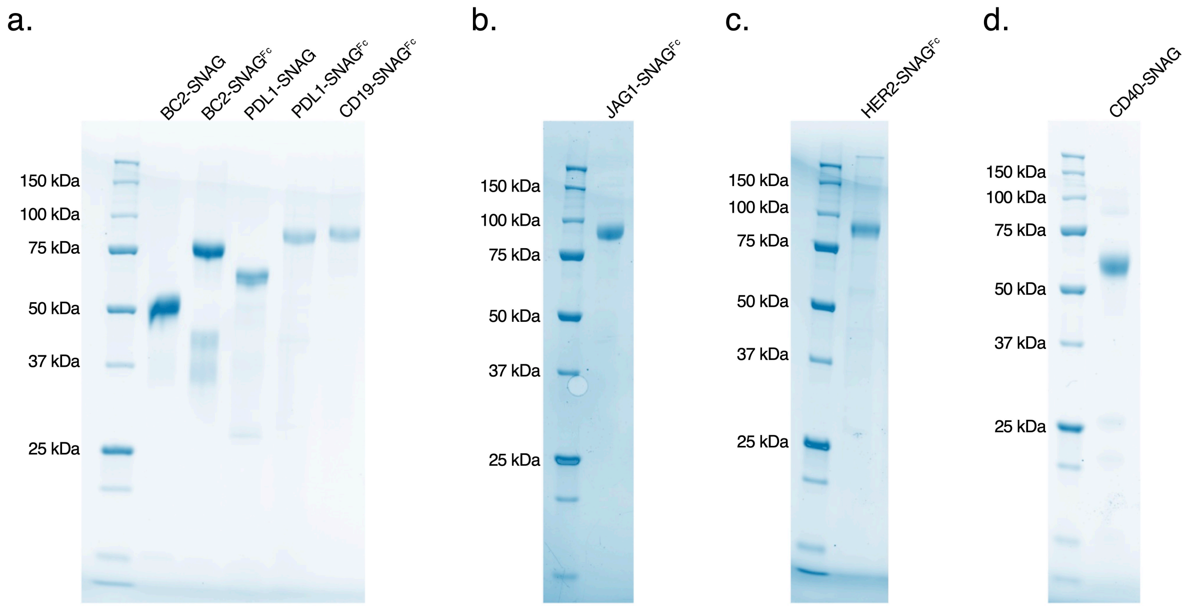


**Figure S2. SDS-PAGE analysis of purified SNAG proteins.** SDS-PAGE was used to evaluate the purity of the indicated SNAGs following nickel and size exclusion chromatography. (a) SDS-PAGE analysis of BC2-SNAGs, PDL1-SNAGs, and the CD19-SNAG. (b) SDS-PAGE analysis of the of the JAG1-SNAG (c) SDS-PAGE analysis of the HER2-SNAG. (d) SDS-PAGE analysis of the CD40-SNAG.

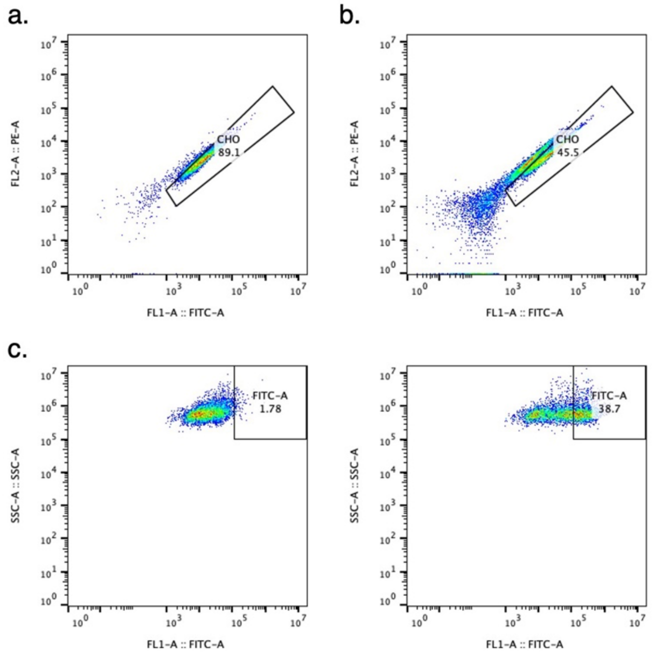


**Figure S3. Gating schemes used to separate Notch1 reporter CHO cells and activated cells.** Flow cytometry gating strategies for selecting Notch1 CHO reporter cells alone (a) or in a coculture with BC2-DLL4$^{HL}$ over-expressing HEK293T cells (b). Gating strategy to determine the population of non-activated and activated Notch1 CHO reporter cells in coculture with BC2-DLL4$^{HL}$ over-expressing HEK293T cells without SNAG (c, left) and with BC2-SNAG$^{Fc}$ (c, right).

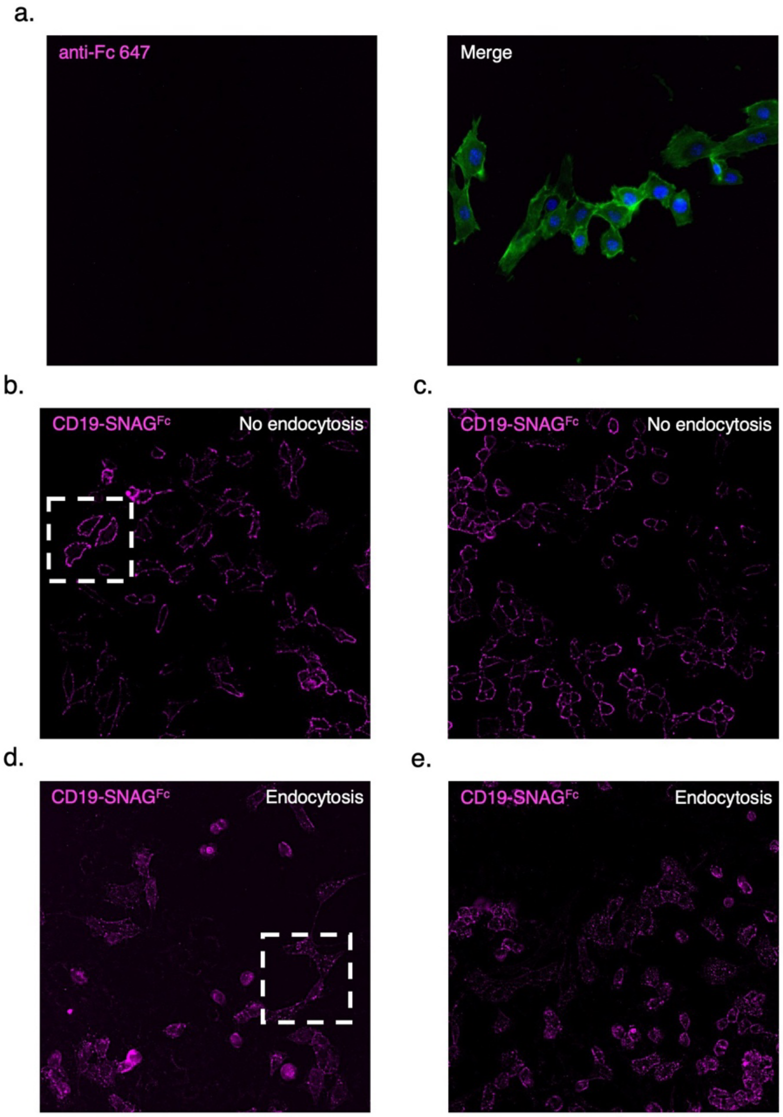


**Figure S4. Immunofluorescent endocytosis assays in 3T3 cells expressing CD19**. (a) Negative control staining utilizing anti-Fc 647 alone (zoom in). Anti-Fc 647 (magenta) left and merged with actin (green) and nuclei (blue) on the right. (b-c) Surface stainings of CD19-SNAG$^{Fc}$-647 (magenta) with multiple cells visualized. The highlighted rectangle in (b) was used as the zoomed-in "no endocytosis" panel for Figure 4a. (d-e) Immunofluorescence images of CD19-SNAG$^{Fc}$-647 (magenta) allowed to endocytose for 15 min with multiple cells visualized. The highlighted rectangle in (d) was used as the zoomed-in "15 min endocytosis" panel for figure 4a.
